# Collective heterogeneity of mitochondrial potential in contact inhibition of proliferation

**DOI:** 10.1101/2023.01.25.525600

**Authors:** Basil Thurakkal, Kishore Hari, Rituraj Marwaha, Sanjay Karki, Mohit K. Jolly, Tamal Das

**Author notes:** All correspondence should be addressed to M.K.J. or T.D.

## Abstract

In the epithelium, cell density and cell proliferation are closely connected to each other through contact inhibition of proliferation (CIP). Depending on cell density, CIP proceeds through three distinct stages, namely the free-growing stage at low density, the pre-epithelial transition stage at medium density, and the post-epithelial transition stage at high density. Previous studies have elucidated how cell morphology, motion, and mechanics vary in these stages. However, it remains unknown whether cellular metabolism also has a density-dependent behavior. By measuring the mitochondrial membrane potential at different cell densities, here we reveal a heterogeneous landscape of metabolism in the epithelium, which appears qualitatively distinct in three stages of CIP. Moreover, it did not follow the trend of other CIP-associated parameters, which increase or decrease monotonically with increasing cell density. Importantly, epithelial cells established a collective heterogeneity in mitochondrial potential exclusively in the pre-epithelial transition stage, where the multicellular clusters of high and low-potential cells emerged. However, in the post-epithelial transition stage, the potential field became relatively homogeneous. The collective metabolic heterogeneity in the pre-epithelial transition stage was independent of the mitochondrial content and spatially correlated with the local cell density. Next, to study the underlying dynamics, we constructed a system-biological model, which predicted the role of cell proliferation in metabolic potential towards establishing collective heterogeneity. Further experiments revealed that the metabolic pattern indeed spatially correlated with the proliferative capacity of cells, as measured by the nuclear localization of a pro-proliferation protein, YAP. Finally, experiments perturbing the actomyosin contractility revealed that while metabolic heterogeneity was maintained in absence of actomyosin contractility, its *ab initio* emergence depended on the latter. Taken together, our results revealed a density-dependent collective heterogeneity in the metabolic field of a pre-epithelial transition stage epithelial monolayer, which may have significant implications for epithelial form and function.

**STATEMENT OF SIGNIFICANCE:** Epithelial contact inhibition of proliferation (CIP) plays a key role in tissue homeostasis, morphogenesis, and development. The biochemical changes in cells during different stages of CIP are not as well-documented as the biophysical changes. We unveil a heterogeneous landscape of metabolism which appears distinct in different stages of CIP. Importantly, in the pre-epithelial transition stage, the epithelial cells establish a collective metabolic heterogeneity wherein multicellular clusters of high and low-potential cells emerge, despite the uniform genetic and nutrient conditions for the cells. The collective heterogeneity is correlated to the local fluctuations in geometrical parameters and the proliferative capacity of cells. Finally, we demonstrate the role of cell mechanics in the establishment of collective heterogeneity.

## INTRODUCTION

The orchestration of cell proliferation and growth in the epithelial tissue is pivotal for morphogenesis, tissue homeostasis, and development in multicellular organisms [1]. This feat is achieved by strict control over the processes like cell division, apoptosis, cell migration, and metabolism. An important way of regulating cell proliferation in the epithelium is contact inhibition of proliferation (CIP). CIP is defined as the decrease in the mitotic rate of the cells when a critical density is attained [2, 3]. On reaching this critical density, the cell-cell junctions and the intracellular molecular signaling undergo significant changes, which prevent the cells to enter the cell division cycle. Importantly, CIP also marks the transition from the free-growing motile state to the highly dense immobile state of the epithelial cells [4]. Loss of CIP can result in cancers and abnormal morphogenesis [5]. Relevantly, depending on mitotic rate patterns, cell shape, and cell density, CIP proceeds through three distinct stages [4]. At low density, cells continue to divide without any decrease in cell mitotic rate. Epithelial cells are observed in a stretched spindle-shaped morphology at low density. At medium density, the cells undergo a morphological transition acquiring a polygonal epithelial shape. Finally, at the highest density, the mitotic arrest happens, and the cell area continues to decrease and induce a kinetic arrest [4, 6]. We refer to these three distinct stages as stages 1, 2, and 3 respectively, henceforth. Relevantly, while some of the biophysical features of these stages of CIP are known, they remain mostly uncharacterized from a biochemical point of view.

Nevertheless, CIP is known to be the cause and result of mechanical and biochemical changes in the epithelial cells [7–10]. As the density increases, stronger cadherin-mediated cell-cell contacts are made, resulting in mechanical changes such as decreasing traction forces [7–9]. These changes eventually act as inhibitory signals resulting in a mitotic arrest [4, 11–14]. The increase in density and contact inhibition also is accompanied by a glass-like phase transition in epithelial cells [15]. At a cell-autonomous level, the mechanism behind the mechanical signaling of CIP is linked to the Hippo signaling pathway. It is known that cell size and shape deformations trigger the Hippo pathway to cause CIP [6, 14, 16–18]. On the other hand, epithelial growth factor (EGF) level is one of the main biochemical cues known for regulating CIP since cadherin-mediated contacts inhibit cell proliferation, only when EGF is below a critical threshold level [10]. Taken together, most of the studies so far have focused on the mechanisms of CIP or biophysical changes associated with CIP. However, metabolic changes that the cells undergo while approaching CIP remain elusive. Previous studies have explored the global metabolic changes of cells as cell density increases. The population-averaged studies - focusing on metabolic changes with increasing density - have revealed that metabolic pointers like oxygen consumption level, net lactate production, ATP content, and total NAD content per cell decrease with increasing cell density [19]. However, the spatiotemporal changes in cell metabolism as the CIP progresses through the aforementioned three stages, remain unknown. Here we investigated the cellular- and multicellular-scale dynamics of metabolism accompanied by different stages of CIP. We asked how the metabolism might be correlated with the geometrical and mechanical parameters of the epithelial cells in the three stages of CIP.

## RESULTS

### Metabolism shows different collectivity at different stages of CIP

To study the spatiotemporal variation of metabolism associated with CIP, we grew confluent MDCK (Madin-Darby Canine Kidney) cell monolayers in varying densities falling in either of three stages. We selected these density values according to the values reported in the previous studies [4, 15] (Fig. 1a). Cell density range 2000-2600 cells/mm^2^ represented stage 1, which corresponded to the free-flowing stage of CIP. Cells at this stage were elongated with many cells showing stretched triangular or quadrilateral shapes (Fig. 1a), and a significant fraction turned out to be in the S-phase as stained by EdU (Supp. Fig. 1a). Next, the density range 2700-4000 cells/mm^2^ represents stage 2, which corresponded to the pre-epithelial transition stage of CIP. Cells at this stage showed cobblestone-like hexagonal or higher-order polygonal shapes but still, a significant fraction turned out to be EdU-positive (Fig. 1a and Supp. Fig. 1a). Finally, in the density range 4400-6000 cells/mm^2^ the fraction of EdU-positive cells decreased with increasing density (Supp. Fig. 1a). This density range represented the post-epithelial transition stage of CIP. Importantly, to obtain monolayers with varying densities but at the same time, to ensure similar culture conditions, we seeded the cells in different initial densities and always imaged them within 24-30 hours post-seeding. As an indicator for the metabolism, especially for oxidative phosphorylation leading to ATP production, we measured the mitochondrial membrane potential (ΔΨ_M_) using tetramethylrhodamine methyl ester perchlorate (TMRM), a cell-permeant fluorescent dye. Since the accumulation of TMRM in the mitochondrial matrix is directly proportional to ΔΨ_M_ [20], we can indirectly read out ΔΨ_M_ using fluorescence microscopy. The signal is bright for polarized mitochondria with high ΔΨ_M_ and is dim if the mitochondrial membrane is depolarized. Despite being genetically identical and maintained under the same environment, we discovered different patterns of metabolic variability emerging at the multicellular level. Further, these variabilities showed different spatial patterns at different stages of CIP, as marked by the cell density (Fig. 1a). In our study, we only considered epithelia post-confluence (i.e., packing fraction is one). Subsequently, at Stage 1, which corresponded to the free-flowing stage of CIP, cells showed cell-cell variability in ΔΨ_M_ without any cluster formation (Fig. 1a, *Left panels*). At stage 2, which corresponded to the pre-epithelial transition stage of CIP, we noticed the emergence of cluster formation, neighboring cells collectively having similar ΔΨ_M_ (Fig. 1a, *Middle panels,* Supp. Fig 1b). Given the collective nature of the metabolic variability in this regime, we termed this observation as collective heterogeneity. Finally, ΔΨ_M_ appeared homogenized at stage 3, which corresponded to the post-epithelial transition stage of CIP (Fig. 1a, *Right panels*). The higher variability of ΔΨ_M_ in stage 2 is confirmed by the higher variance compared to that of stages 1 and 3 (Fig. 1b). At the same time, the TMRM intensity distribution shifted from a nearly Gaussian distribution at stage 1 to a long-tail distribution at stage 2. At the same time, a significant fraction of the cell population showed vanishingly small ΔΨ_M_ at stage 2 (Fig. 1c and Supp. Fig. 1b, red arrows). This population was unique to stage 2 and disappeared again at stage 3. At stage 3, a skewed Gaussian distribution reappeared (Supp. Fig. 1b), implying that ΔΨ_M_ of neighboring cells lost their correlation at this stage. This result is surprising given that the correlation length of cellular motions and forces monotonically increases with increasing density [15, 21].

**Figure 1:**
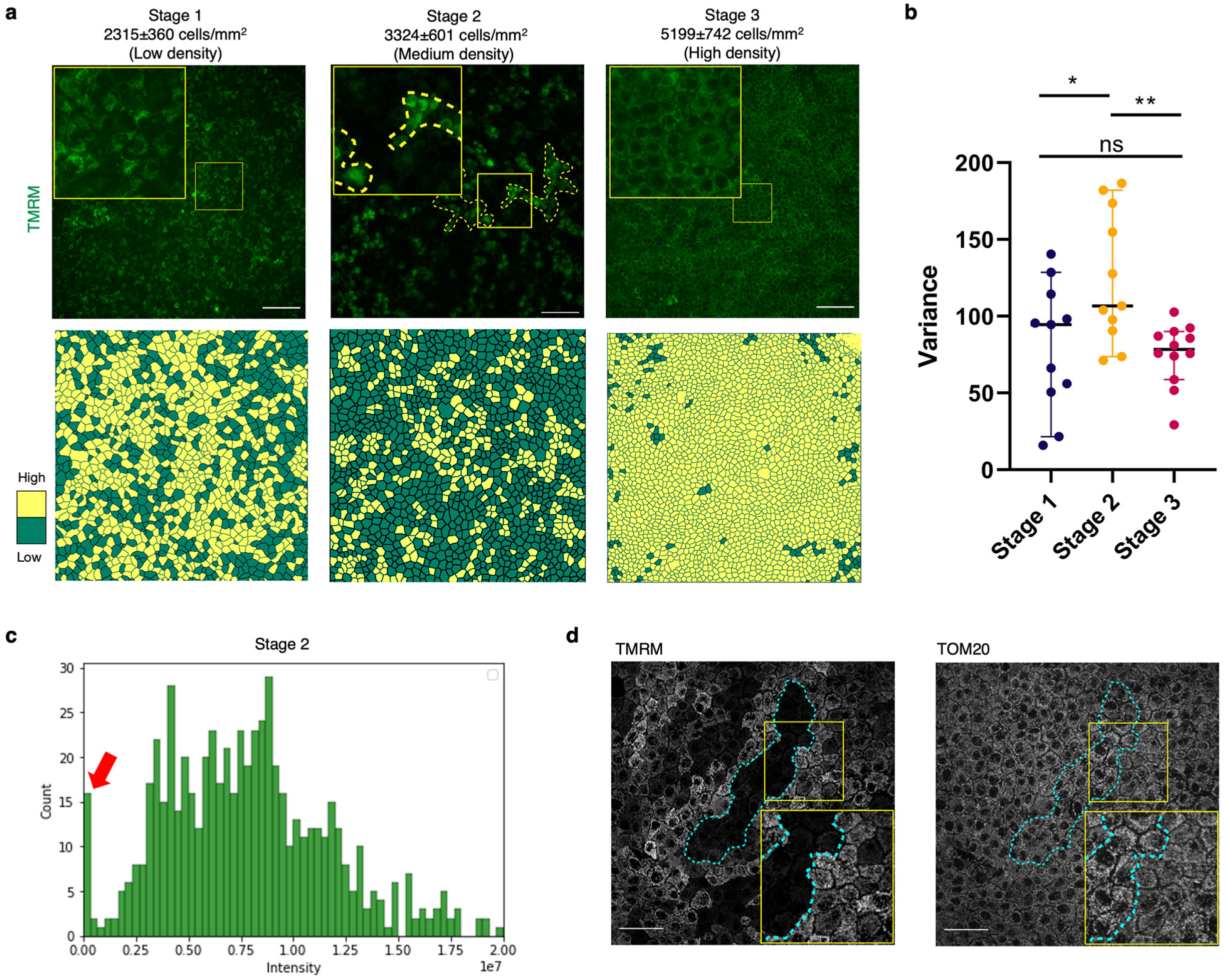
Metabolism shows different patterns of collectivity at different stages of CIP. **a)** *Top panel:* TMRM stained images of confluent MDCK monolayer corresponding to the three global density regimes, showing differences in the metabolic heterogeneity pattern. Left to right: Low, medium and high density. Yellow dashed lines mark the clusters formed in medium density. Scale bar 100 μm. The mean and standard deviation of the number density corresponding to each stage are mentioned on top of the panels. *Bottom panel:* Cartoon of correlation length variation at different global densities corresponding to the FOV in Fig. 1a. Yellow cells show high ΔΨ_M_ cells and green square shows low ΔΨ_M_. 25% of the maximum normalized intensity is considered as the threshold for low and high classification here. **b)** Box and whiskers plot of variance of TMRM intensity for different density regimes in MDCK monolayer. Statistical significance was assessed using an unpaired Student’s t-test with Welch’s correction (two-tailed). **c)** Representative histogram showing the distribution of TMRM intensity for cells in stage 2 of CIP. The peak with near-zero intensities is marked by the red arrow. **d)** Confocal images of TMRM-stained MDCK cells in the left panel and right panel show the Tom20 immunofluorescence confocal image. Scale bar 50 μm.

To check whether the metabolic variability was TMRM-specific, in separate samples, we stained the cells with another ΔΨ_M_ indicator dye JC-1, where the measurement becomes ratiometric. JC-1 exhibits ΔΨ_M_-dependent accumulation in mitochondria, indicated by a fluorescence emission shift from green (~525 nm) to red (~590 nm), as dye aggregates are formed from the monomers on the mitochondrial membranes. Hence red:green intensity ratio is high for polarized mitochondria and low for depolarized mitochondria. Moreover, the JC-1 signal is known to be independent of the morphological parameters of mitochondria. JC-1 staining also captured the different patterns of metabolic variability at different stages (Supp. Fig 1d). Like TMRM staining, we observed collective heterogeneity in JC-1 red emission at stage 2, and more homogenous signal at stages 1 and 3 (Supp. Fig 1d). Hence, the results from JC-1 staining experiments indicated the observed density-dependent heterogeneity of the ΔΨ_M_ was independent of the choice of dye. Finally, we investigated whether differences in ΔΨ_M_ stem from the cell-cell differences in the mitochondrial mass. For this, we first stained cells at stage 2 with TMRM and imaged them. TMRM intensity appeared heterogeneous here. Later, we fixed the same sample and stained it with an antibody against a mitochondrial outer membrane protein, Tom20, that is commonly used to quantify the mitochondrial content [22]. Tom20 signal was homogenous in the field of views where TMRM was heterogeneous (Fig. 1d). Even the cells with negligible ΔΨ_M_ have normal mitochondrial content. The result suggests that the drop in ΔΨ_M_ is not due to the absence of mitochondria content, but rather a functional change. Taken together, these results revealed a heterogeneous ΔΨ_M_ field emerging in the epithelial monolayer, whose length scale and variance depended on the global cell density. This heterogeneity was both qualitatively and quantitatively different in three previously reported stages of CIP. Importantly, these results revealed an emergence of collective heterogeneity at stage 2, which is also the pre-epithelial transition stage. Hence, going forward, we investigated how this collective heterogeneity observed at Stage 2 depended on the local parameters, beyond its dependency on the global density.

### Correlation of local variations in membrane potential with the local geometric environment

Relevantly, at stage 2, cells display a local fluctuation in density across the monolayer field [23, 24]. Hence, we next studied how the local fluctuation in density influences the metabolism in the pre-epithelial transition stage. For this purpose, we mapped the collective heterogeneity in metabolism, as measured by the spatial variation in TMRM intensity, onto the local number density of cells. To this end, we segmented the cell boundary from brightfield images using a custom-written code in MATLAB. The local density value was defined, by counting the number of cells around each pixel in a unit neighborhood area. The unit neighborhood area was defined as, a square with a side of 3 times the average cell diameter. Because a side size of 3 times the average cell diameter was the optimum neighborhood-area to give the maximum difference between average ΔΨ_M_ for low and high local density population of cells. (Supp. Fig 2a-b). Further correlation analysis showed that ΔΨ_M_ was correlated with local variation of the number density of cells (Fig. 2a, Supp. Fig 2b). To evaluate the statistical significance of this observed correlation, we compared the TMRM intensity of the top 20% dense region (high density) with that of the bottom 20% dense region (low density) (Fig. 2b). The comparison revealed that TMRM intensity in high-density regions was higher than TMRM intensity in low-density regions (Fig. 2b). Relevantly, local fluctuations in local density can result in changes in the other geometrical parameters of cells as well. We, therefore, investigated how different geometric parameters, including the cell area, cellular aspect ratio, and shape index are spatially correlated with TMRM intensity. Aspect ratio is defined as the ratio of the minor axis and major axis of the cell. The shape index is defined as the ratio of the perimeter to the square root of the area. We considered the cells of lower quartile TMRM intensity and upper quartile TMRM intensity as the cells with low and high ΔΨ_M_ for quantification purposes. The results delineating the variation of TMRM intensity with cell area revealed that the cells with high ΔΨ_M_ were smaller compared to the cells with lower ΔΨ_M_ (Fig 2c left panel). At the same time, the results delineating the variation of TMRM intensity with cellular aspect ratio and shape index revealed that the cells with high ΔΨ_M_ were more rounded compared to cells with low ΔΨ_M_, which were extended. (Fig. 2c middle and right panel). To evaluate how the mitochondrial content is different in the high and low ΔΨ_m_ populations, we quantified the total Tom20 intensity in both populations. Cell population with high ΔΨ_M_ has relatively higher mitochondrial content and vice versa (Supp. Fig 2c). Cells reducing in size in high-density regions may cause the mitochondria to be tightly packed, resulting in a higher value of ΔΨ_M_ when stained. Counterintuitively, we argue that it is not the case, because (i) there is high contrast in ΔΨ_M_ between the population in higher resolution images (Supp. Fig 1b) (ii) Tom20 signals look homogenous which shouldn’t be the case if the mitochondrial crowding due to cell size variation has a major effect on the imaging. Taken together, these results show that beyond the global density, locally the heterogeneity in ΔΨ_M_ is connected to the heterogeneity in geometric factors but not to the biochemical factor such as mitochondrial biogenesis.^9^

**Figure 2:**
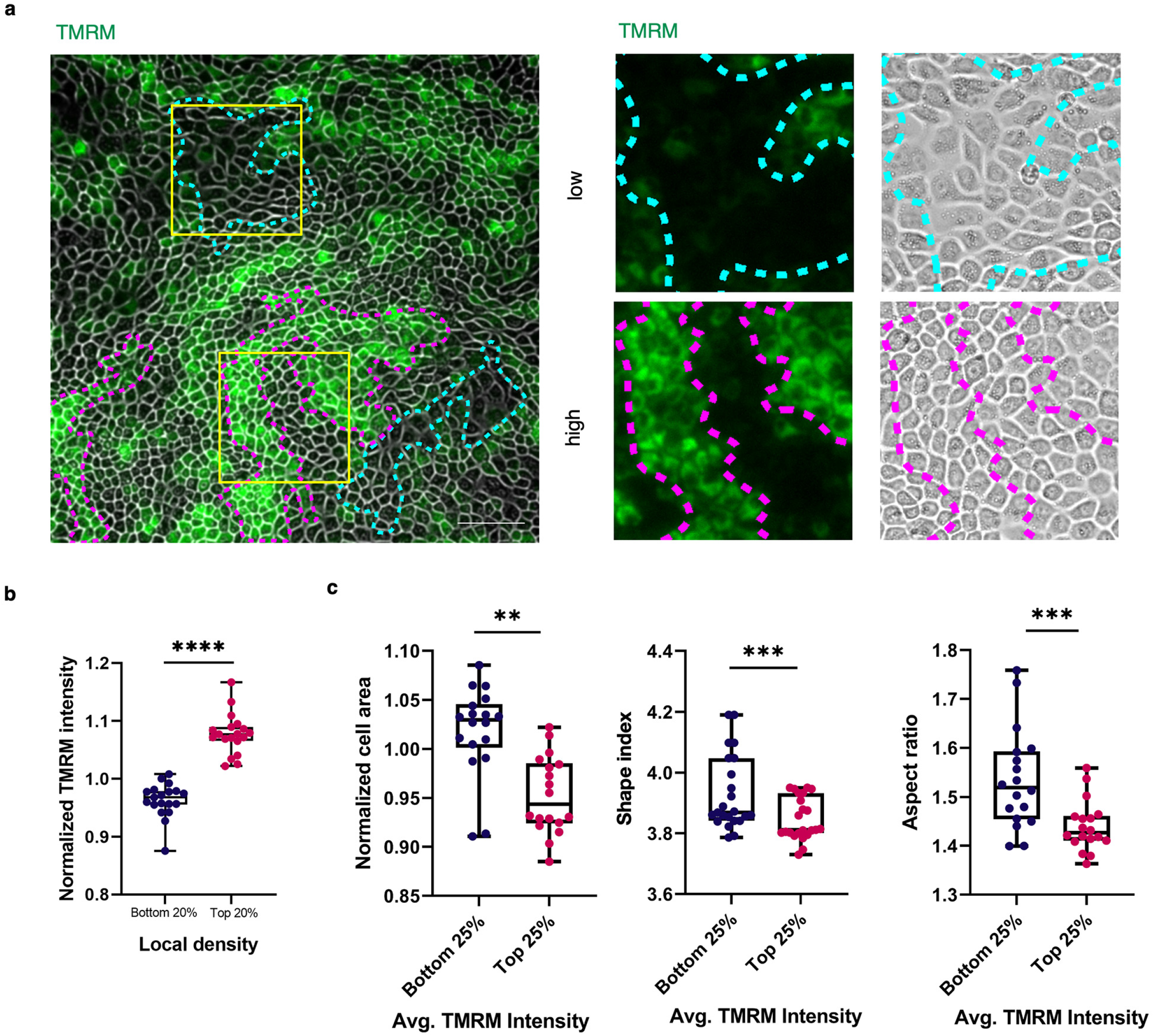
Local variations in membrane potential is correlated with the local geometric environment. **a) Left:** Images of TMRM stained MDCK cells, merged with the bright field image showing cell boundaries. The increase and decrease in TMRM intensity correlate with the changes in local density. Magenta dashed lines mark high ΔΨ_M_ and cyan dashed lines mark the low ΔΨ_M_ cells. Scale bar 100 μm. **Right:** Insets showing the high and low density corresponding to the changes in TMRM intensity. The yellow square in the left panel shows the cells used for insets. **b)** Box and whiskers plot showing the difference in mean TMRM intensity for the MDCK cells at the highest (top 20%) and lowest (bottom 20%) local density neighborhood. Statistical significance was assessed using Wilcoxon matched-pairs signed rank test **c) Left:** Box and whiskers plot showing the difference in the mean area for the MDCK cells with highest (upper quartile) and lowest (lower quartile) TMRM intensity **Middle:** Box and whiskers plot showing the difference in mean shape index for the MDCK cells with highest (upper quartile) and lowest (lower quartile) TMRM intensity **Right:** Box and whiskers plot showing the difference in mean aspect ratio for the MDCK cells with highest (upper quartile) and lowest (lower quartile) TMRM intensity. Statistical significance was assessed using Wilcoxon matched-pairs signed rank test.

### Dynamical model

We next asked how the metabolic heterogeneity is established as a function of cell proliferation. To this end, we constructed a phenomenological population-level dynamical model. Based on the results of dependence of metabolic heterogeneity on density and CIP, we make two assumptions in the model: a) Cell proliferation decreases (therefore the proliferation time increases) sigmoidally with local cell density, and b) ΔΨ_M_ termed “activation” in the model, sigmoidally increases with local cell density. The sigmoidal dependence was chosen due to two reasons: a) both the proliferative capacity and activation threshold have an upper and lower bound, and b) it allows quantifying two key parameters: threshold local densities that mark a steep transition from high to low proliferative capacity (proliferation threshold) and that from a low to high ΔΨ_M_ (activation threshold). These thresholds can potentially be experimentally determined using temporally collected images of cell culture. One crucial difference between the model and experimental condition is that the model does not operate at confluent conditions, but confluence is the end goal of the model. This difference can be justified by the fact that the key assumptions are not dependent on the confluence but on local cell density, which can be captured by the measure of density defined in the model as the number of cells in a 5-cell square.

Our model was able to capture the three stages of CIP-linked metabolic heterogeneity observed experimentally: a sporadic activation at low density (Stage 1), patches of activation representing collective heterogeneity (Stage 2) and homogeneous activation (Stage 3), as shown in Fig 3a. Stage 1 is characterized by a narrow distribution of activation with low mean. Stage 2 is characterized by the activation distribution skewed to left, with patches of activation in the field as mentioned above. Stage 3 is characterized by a narrow distribution with high mean activation (Supp. Fig. 3a). Given the three stages, we wanted to identify factors that control the dynamics of these stages, specifically the residence time in each of the stages. As a first step, we focused on qualitative predictions that can be verified experimentally. First, we tried to understand the appearance of collective heterogeneity as a function of the threshold of activation and threshold of proliferation. Given the distribution characteristics above, we tracked the dynamics of the variance of cell activation level distribution with time as a metric to distinguish between the three stages (Fig 3b, Supp. Video 1a). Our initial simulation parameters had a high activation threshold (0.9) and low proliferation threshold (0.1). We first fixed the activation threshold and varied the division threshold (Fig 3c, Supp. Video 1b). We found that having a low proliferation threshold in this scenario allows for a clear emergence and longer sustenance of stage 2, where collective heterogeneity exists. At higher division thresholds, stage 2 emerges, but remains for a shorter period of time. We then repeated the experiment at a lower activation threshold and found multiple, short-lived patches of high variance activation profiles (Fig 3d, Supp. Video 1c) for each division time. As the density is constantly increasing, the dependence of the emergence of high variance patches on the density is lost, along with the stable collective heterogeneity observed in the previous scenario. Therefore, we infer that a high activation threshold is enough to enable the existence of collective heterogeneity, while a low division threshold stabilizes the collective heterogeneity. Furthermore, we predict that the dependence of activation and proliferation must be on local density as opposed to global density, as the dependence of ΔΨ_M_ on global density leads to rapid homogenization of the ΔΨ_M_, thereby eliminating the existence of collective heterogeneity. The neighbourhood parameter, R, in our simulation determines whether a cell is sensitive to local density (low value of R) or to high density (high value of R). We find that we the value of R increases, the variance in activation level that distinguishes collective heterogeneity goes down, being comparable to the variance of low- and high-density scenarios at R=9 (Fig 3e). Taken together, the simulation results predicted a role of cell proliferation in metabolic potential towards setting the collective heterogeneity in the latter at medium density.

**Figure 3:**
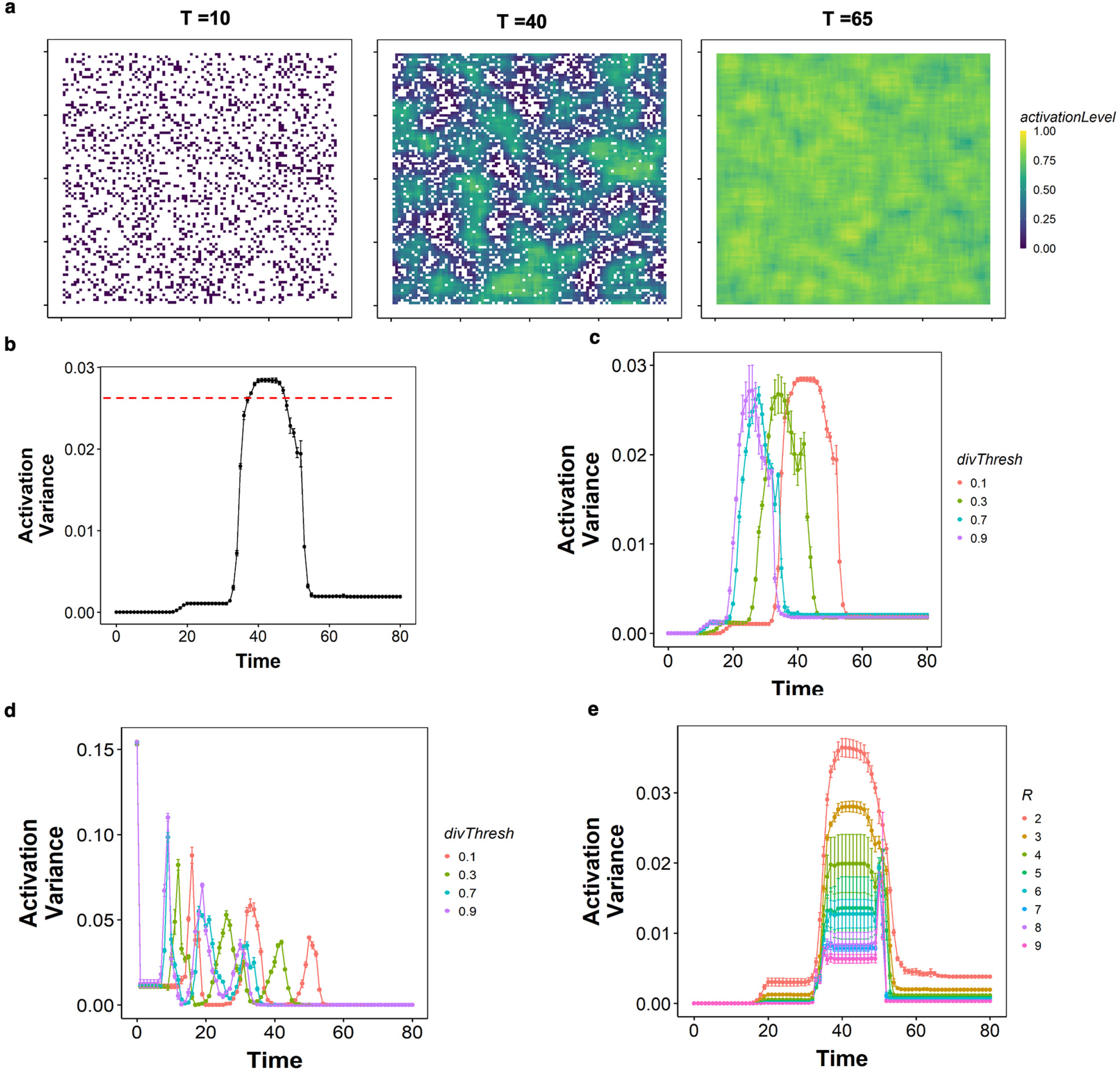
Results from the simulation. **a)** Activation level heatmaps at different time points (T=10,40,65) corresponding to 3 stages **b)** Line plot showing activation variance against time points (each timepoint corresponds to different densities) **c)** Lineplot showing activation variance for different division threshold values with high activation threshold (0.9)**. d)** Same as c, but for a low activation threshold (0.1). **e)** Lineplot showing activation variance against the neighborhood radius. Maximum variance is obtained at 2-3 cell neighborhood radius.

### Metabolism pattern emergence is correlated to the proliferative capacity of cells

Next, to test this prediction from the dynamical model, we studied ΔΨ_M_ and the proliferative capacity of cells in the same sample. As a proxy for proliferative capacity, we used nuclear/cytoplasmic (N/C) localization of a transcriptional regulator protein, YAP (yes-associated protein 1). Relevantly, we avoided EdU staining here we reasoned that EdU-positive S-phase cells would be timewise advanced in the cell cycle. In that case, even if a change in the metabolic potential had initiated the proliferation earlier, S-phase cells might not be correlated to the spatial distribution of metabolic patterns at the time of observation. On the other hand, YAP is a protein that when localized to the nucleus induces division by activating genes involved in proliferation and suppressing those involved in apoptosis. Nuclear localized YAP suggests a proliferative cell and cytoplasmic localization suggests inhibited proliferation [18]. YAP nuclear localization, therefore, indicates a very early stage of pro-proliferation activity, perhaps right at the moment when the change in metabolism triggers proliferation. First to understand how the proliferative capacity of cells changes collectively at different stages of CIP, we immuno-stained confluent MDCK cells with a YAP-antibody at different densities. At Stage 1, all cells have nuclear localization of YAP. At stage 2, cells appear as patches in terms of YAP N/C localization like the collective heterogeneity in metabolism. At Stage 3, all cells have cytoplasmic localization of YAP (Suppl. Fig. 4a). The results indicate the variability of proliferative capacity among cells is similar to that of ΔΨ_M_, especially the appearance of collective patches in Stage 2. Hence, we further studied the local correlation of proliferative capacity with ΔΨ_M_ at Stage 2. To this end, we stained the cells at Stage 2 with TMRM to measure ΔΨ_M_. We then fixed and immunostained the same cells with YAP to find that, YAP N/C localization was correlated with the ΔΨ_M_. The low ΔΨ_M_ regions were associated with the cells having predominantly nuclear YAP localization, while the high ΔΨ_M_ regions were associated with the cells having predominantly cytoplasmic YAP localization (Figs. 4a-b). For the quantification, we considered the cells of lower quartile TMRM intensity and upper quartile TMRM intensity as the cells with low and high ΔΨ_M_. The results delineating the variation of TMRM intensity with YAP nuclear translocation revealed that the cells with low ΔΨ_M_ have proliferative capacity while the cells with higher ΔΨ_M_ are entering quiescence. Cells with low ΔΨ_M_ have high nuclear translocation of YAP compared with the cells with high ΔΨ_M_ (Fig 4c). Together these results upheld the predicted correlation between proliferative capacity, as represented by the nuclear localization of YAP, and the metabolic heterogeneity, suggesting a connection between these two aspects of a confluent epithelial monolayer.

**Figure 4:**
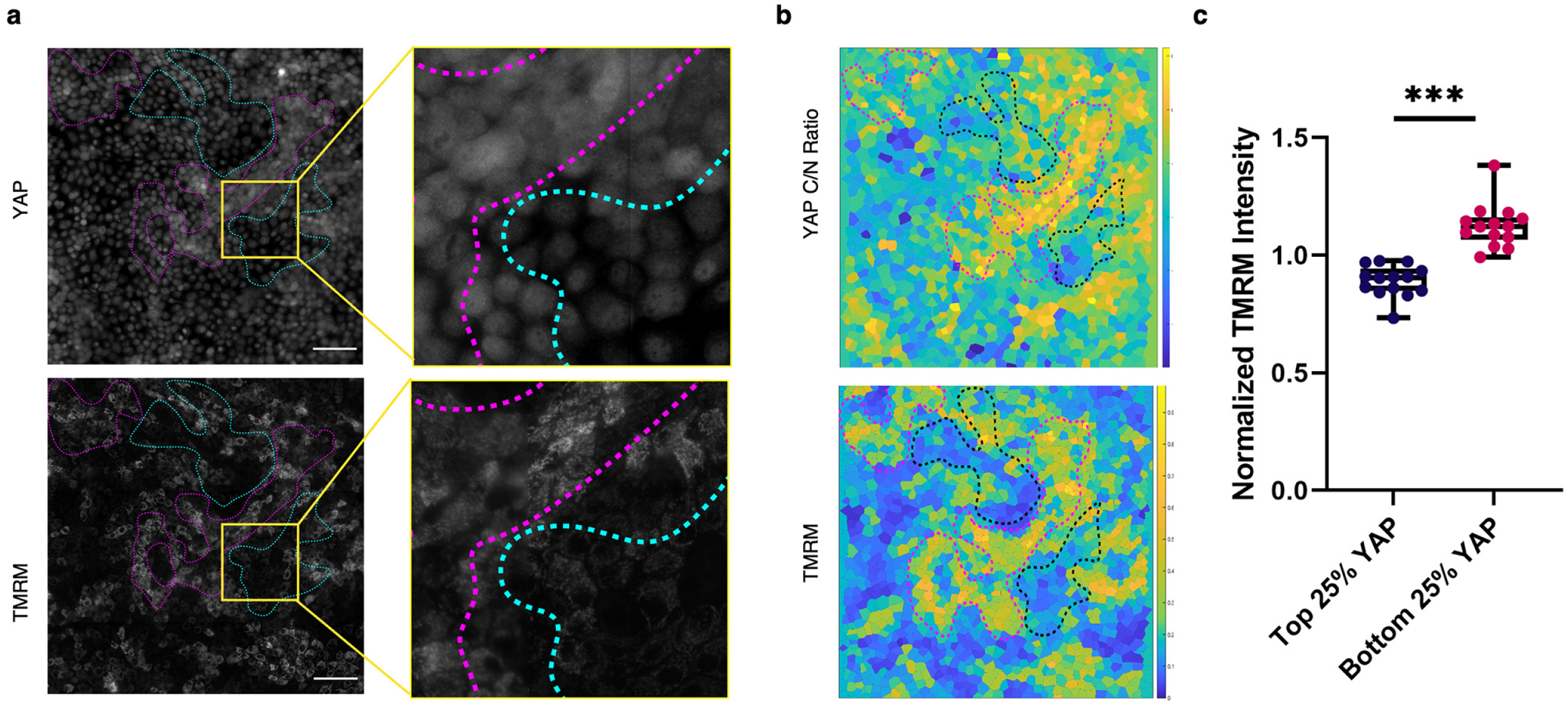
Metabolic heterogeneity is correlated to the proliferative capacity of the cells. **a)** Fluorescence images comparing YAP nuclear/cytoplasmic localization (top panel) and TMRM levels (bottom panel) in MDCK cells at medium global density regime. Nuclear localization of YAP corresponding with low TMRM intensity is marked with cyan dashed lines. Cytoplasmic localization of YAP corresponding with high TMRM intensity is marked with magenta dashed lines. On the right, insets are shown at higher magnification. Scale bars are 100 μm. **b)** Heat maps showing mean TMRM levels (bottom panel) and YAP cytoplasmic/nuclear ratio (top panel) in MDCK cells. Dashed lines show the cell population with high and low TMRM intensity and YAP C/N ratio. **c)** Box and whiskers plot showing the difference in mean TMRM intensity in cell population having lowest (lower quartile) and highest (upper quartile) nuclear/cytoplasmic ratio of YAP. Statistical significance was assessed using the Wilcoxon matched-pairs signed rank test.

### Metabolic memory and dependence of collective heterogeneity on the active mechanical state

To summarize the results obtained so far, we showed that the collective heterogeneity in metabolism is associated with local density fluctuations and proliferation. Notably, both local density fluctuations and proliferation are mechanosensitive events. With local density fluctuation of the epithelial monolayer, actomyosin content and organization - key players in cell mechanics - are known to change [25, 26]. The proliferation regulator YAP/TAZ is also mechanosensitive [26]. In addition, contact inhibition of proliferation ceases to happen in the absence of adhesion molecules or in actomyosin-inhibited conditions [27, 28]. Lastly, mechanical stretch can increase the proliferation, acting through the Hippo pathway [27]. Hence, we next asked whether actomyosin contractility is necessary to maintain and/or establish the metabolic collective heterogeneity. First, to check whether cells can maintain the collective heterogeneity in absence of actomyosin contractility, we treated the cell monolayer at Stage 2 with a myosin inhibitor, blebbistatin, that is known to reduce actomyosin contractility. After the treatment, we tracked the cells for 4 hours but did not observe any change in metabolic heterogeneity (Fig. 5a). Similarly, treatment of the cells with Y27632, a ROCK inhibitor (Rho-associated, coiled-coil containing protein kinase) did not cause any change in metabolic heterogeneity over time (Fig. 5b). This experiment indicated that the maintenance of collective heterogeneity does not depend on active cellular mechanics. Second, to check whether the establishment of collective heterogeneity requires actomyosin contractility, we first abolished the ΔΨ_M_ in all cells by treating the cells with an H^+^ ionophore and uncoupler of oxidative phosphorylation, FCCP, and then studied how the cells regained the ΔΨ_M_ upon withdrawal of FCCP from the medium, in both presence and absence of blebbistatin and Y27632. As expected, FCCP treatment homogenously depolarized the mitochondria of all cells in the monolayer (Fig 5c). When FCCP was washed off post-treatment and normal DMEM media was added, the cells re-established the same pattern of collective heterogeneity that they had before depolarization (Fig. 5c, top panel). However, when the recovery DMEM contained blebbistatin, the cells regained their ΔΨ_M_, but the collective heterogeneity of ΔΨ_M_ vanished and rather a metabolically homogenous population emerged (Fig. 5c, middle panel). Whereas, when the recovery DMEM contained Y27632, the cells regained their ΔΨ_M_, but with a visibly different pattern of collective heterogeneity of ΔΨ_M_. In contrast to the blebbistatin treatment, ROCK inhibition did not homogenize the ΔΨ_M_ (Fig. 5c, bottom panel). These results revealed that once emerged, epithelial cells can maintain the metabolic heterogeneity in absence of actomyosin contractility. However, its initial emergence depended on the latter. Finally, we investigated whether there exists a direct correlation between the local mechanical state and the collective metabolic heterogeneity. To this end, we calculated the cell-cell junctional tension and pressure using the Bayesian force inference method [29] and at the same time, stained the cells with TMRM (Supp Fig 5a). For the quantification, we considered the cells of lower quartile TMRM intensity and upper quartile TMRM intensity as the cells with low and high ΔΨ_M_. The results delineating the variation of TMRM intensity with cell pressure revealed that the cells with high ΔΨ_M_ have higher intracellular pressure compared to the cells with lower ΔΨ_M_ (Fig. 5d). Together these results indicate the importance of active cellular mechanics on the establishment of collective heterogeneity in metabolism. However, maintenance of the metabolic variability does not depend on actomyosin contractility, only the *ab initio* emergence does.

**Figure 5:**
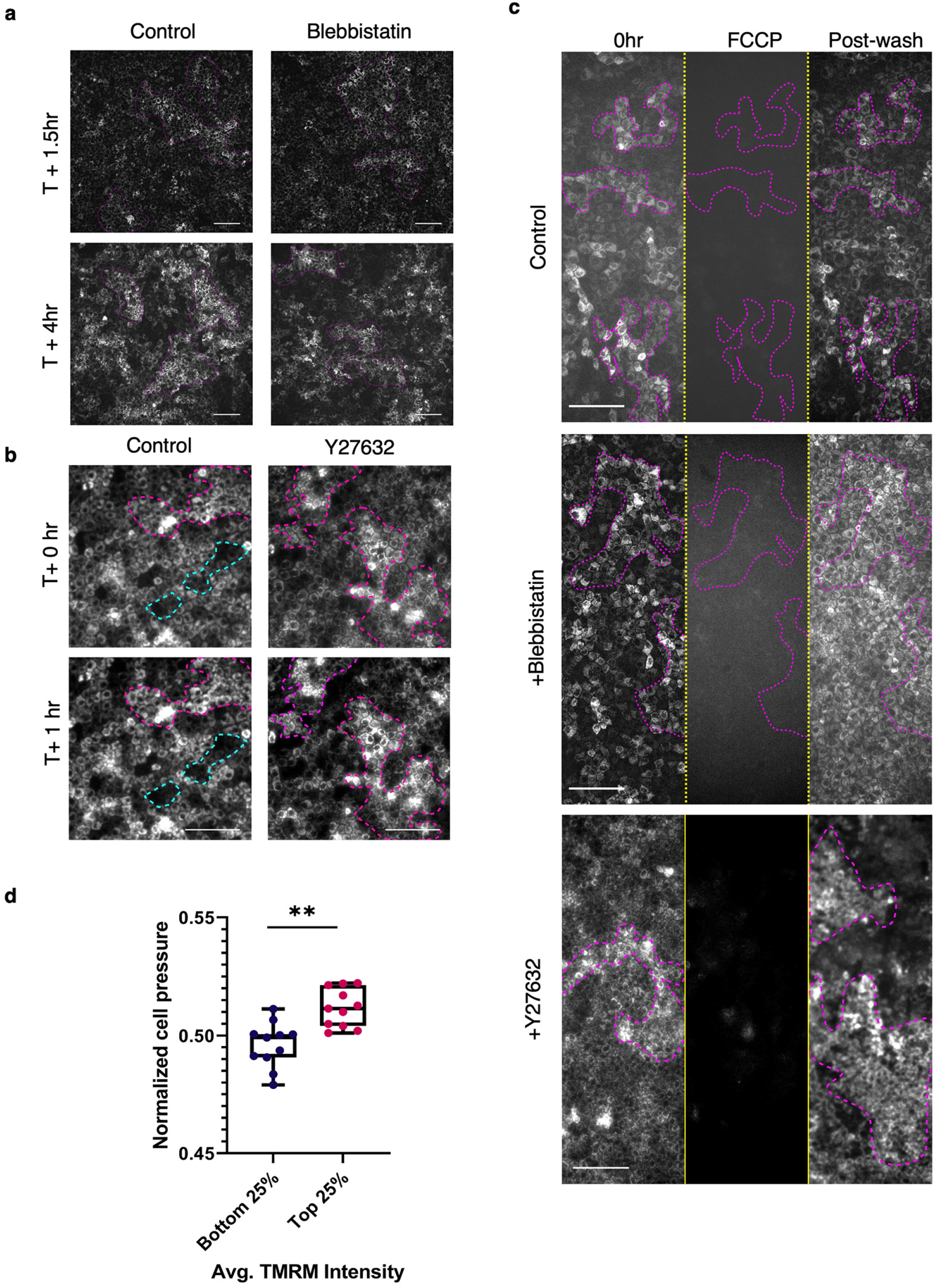
Metabolic memory and dependence of collective heterogeneity on active mechanical state. **a)** TMRM stained images of MDCK cells monolayer at stage 2 showing patterns of TMRM heterogeneity. Treatment with blebbistatin without FCCP did not get rid of metabolic heterogeneity. *Left panels:* control, *right panels:* blebbistatin treated. Scale bar 100μm. **b)** TMRM stained images of MDCK cells monolayer at stage 2 of CIP. Treatment with Y27632 without FCCP did not get rid of metabolic heterogeneity. *Left panels:* control, *right panels:* Y27632 treated. Scale bar 50μm. **c)** A time-lapse montage of TMRM stained MDCK monolayer at stage 2, left to right: before FCCP treatment, after FCCP treatment and after washing off FCCP with replacement of complete DMEM media. *Top panel:* vehicle wash-off media could bring back the initial metabolic heterogeneity patterns in the monolayer. *Middle panel:* wash-off media with blebbistatin was used, heterogeneity was lost in this case post-wash. Scale bar 100um. Collective heterogeneity is highlighted with magenta dashed lines. *Bottom panel:* wash-off media with Y27632 was used, the pattern of heterogeneity changed in this case post-wash. Collective heterogeneity is highlighted with magenta dashed lines. Scale bar 100μm. **c)** Box and whiskers plot showing the difference in mean TMRM intensity in cell population having lowest (bottom 20%) and highest (top 20%) cell pressure. Statistical significance was assessed using the Wilcoxon matched-pairs signed rank test.

## DISCUSSION

While the metabolic changes in a cell population as CIP sets in have been investigated, the patterns of local metabolic changes as the cells progress through CIP are relatively less known. Collectively, our experiments established three different metabolic regimes corresponding to the three different stages of CIP (Fig. 1a). We unveil the emergence of a multicellular-level collective heterogeneity in ΔΨ_M_ at stage 2 of CIP. Importantly, this collective heterogeneity in ΔΨ_M_ was exclusive to stage 2 and disappeared at stage 3, while other previously studied parameters such as the correlation length of cellular motions and the monolayer stresses monotonically always increase with increasing density. Our further investigations show that the metabolic changes are correlated to the local changes in the cell density and the proliferative capacity of cells (Fig 2,4). While the connection between CIP and mechanics has been known previously [4, 30], here we demonstrate that the local metabolic changes that accompany CIP are also connected to cell mechanics (Fig. 5). The system under our study is a monolayer of genetically identical epithelial cell populations - all under the same nutrient conditions. Hence, the self-emergence of a collective-level heterogeneity in ΔΨ_M_, in the absence of any external signaling in an otherwise identical environment, is of great interest. Through the experimental characterization and the systems biology model, we suggest that epithelial cells bring about these biochemical variabilities because of their intricate relations with emerging biophysical differences. Relevantly, the biophysical differences in terms of geometrical and mechanical properties of the cells are basically a result of the non-uniform growth in the monolayer. Non-uniform growth gives rise to feedback mechanical signals to ensure a uniform growth [24]. Hence, we speculate that the metabolic variabilities in cells lead to mechanical variabilities. However, further studies are required for deeper understanding.

Subsequently, we found that the ΔΨ_M_ variabilities were correlated to the proliferative capacity of cells (Fig. 4). Quiescent cells have higher ΔΨ_M_ while proliferating cells have less. It is possible that this is because the proliferating cells prefer the glycolytic pathway rather than OXPHOS for their energy requirements, hence staying at the basal ΔΨ_M_ level, an effect similar to the Warburg effect [31]. Interestingly, similar density-dependent changes have been observed in contact inhibition of locomotion (CIL), in the case of jamming transition in epithelial cells [15, 32]. In jamming-transition, as density increases, the velocity of cells decreases but the fastest cells move in large multicellular groups [15]. Cells become less elongated and less variable, as they get approach jamming [33]. It also is known that during epithelial unjamming, energy metabolism shifts towards the glycolysis [34], which connects the contact inhibition of locomotion (CIL) to local changes in metabolism. In our work, we reveal the local changes in metabolism with CIP. Hence, these results together indicate that density-dependent changes in the cellular metabolism is a fundamental property of the epithelial tissue in the context of both CIP and CIL, which should have enormous physiological consequences.

In this regard, one of the main physiological implications of the density-dependent collective heterogeneity will be in the pattern formation in multicellular organisms. Hair-follicle distribution in mammalian skin is a scenario where this may be relevant. Which cells among the skin give rise to placodes and what the determining factor is still not completely understood [35, 36]. Given the density and shape changes in cells during the placode formation, it will be interesting to look at the metabolic and biophysical changes before the initiation of placode formation. As we used a monolayer of epithelial cells for the current study, fundamental characters of the epithelia in the absence of any external signaling can be pinpointed here. At the same time, this becomes a limitation since it is not the case in physiological conditions. An investigation into the local metabolic changes in a 3D organoid model along with the biophysical characterization will be an interesting study. We have also solely concentrated on ΔΨ_M_ reading as a proxy for metabolism. A complete metabolic profiling as the cells progress through CIP with increasing density will be a worthwhile endeavor, given the results from our current study.

## MATERIALS AND METHODS

### Cell culture

Madin-Darby canine kidney (MDCK) epithelial cell lines were used for this study. Tetracycline-resistant Wild-type MDCK (MDCK-WT) cell lines were a gift from Yasuyuki Fujita. Cells were cultured in Dulbecco’s modified Eagle’s medium supplemented with GlutaMax (Gibco) with 5% fetal bovine serum (tetracycline-free FBS, Takara Bio) and 10 U mL^-1^ penicillin and 10□μg mL^-1^ streptomycin (Pen-Strep, Invitrogen) in an incubator maintained at 37°C and 5% CO2, unless mentioned otherwise. All transient plasmid DNA transfections were done using Lipofectamine 2000 (Invitrogen) following the manufacturer’s protocol. The density of cultured cells we varied by increasing the seeding density. To get higher density with desired control, the cells were seeded in an ibidi 2 well insert.

### Staining for the mitochondrial membrane potential

We used either JC-1 Dye (2μg/mL Invitrogen cat. No. T3168) or TMRM (100 nM, Image-iT TMRM, Cat. No. I34361, Invitrogen) to stain for ΔΨ_M_. The cells were incubated at 37°C for 30 mins with TMRM for staining the cells. Following the incubation, cells were imaged immediately in TMRM containing media. In the case of JC-1 imaging, the dye-containing media is replaced with DMEM media prior imaging.

### Antibodies used

The following antibodies were used for this study. Tom20 (D8T4N) Rabbit mAb (Cell signalling technology #42406, 1:200), YAP/TAZ (D24E4) Rabbit mAb (Cell signalling technology #8418, 1:100). Alexa Fluor-conjugated secondary antibodies were used at the same dilution as that of corresponding primary antibody.

### Immunofluorescence

Cell fixation was done with 4% formaldehyde diluted in 1x phosphate-buffered saline (PBS, pH 7.4) at room temperature (RT) for 10 minutes, followed by 1X PBS washes (three times). Cell permeabilization was carried out with 0.25% (v/v) Triton X-100 (Sigma) in PBS for 10 min at RT followed by washing thrice with PBS to remove the reagent. To block non-specific antibody binding samples were incubated in 2% BSA in PBST (0.1% v/v Triton X-100 in 1X PBS) at RT for 45 minutes. The blocking buffer was removed after 45 minutes, and the primary antibody dilution prepared in the blocking buffer was added to the samples for 60 minutes at RT or at 4°C overnight. Afterwards, samples were washed twice with PBST and thrice with 1X PBS. Next, secondary antibodies tagged with fluorophore were (same dilution as primary) prepared in blocking buffer and added to the sample for 60 minutes at RT. To counterstain cell nuclei, the samples were added with a DNA-binding dye 4’,6-diamidino-2-phenylindole (DAPI, 1 μg mL^-1^ in PBS, Invitrogen) along with the secondary antibody solution. Lastly, thorough washing of the samples was done with PBST and PBS before imaging.

### EdU proliferation assay

We used Click-iT™ Plus EdU Cell Proliferation Kit (Invitrogen C10637) or Click-iT™ EdU Imaging Kit (Invitrogen C10086) was used for the proliferation assay. MDCK cells were seeded to the confluence at the required density, in a glass bottom dish. The cells were incubated at 37°C in 10μM EdU (5-ethynyl-2’-deoxyuridine) diluted in complete DMEM media for 60 minutes. Followed by cell fixation with 4% formaldehyde diluted in 1x phosphate-buffered saline (PBS, pH 7.4) at RT for 15 minutes. Fixation media was removed, and the cells were washed twice with 3% BSA solution in PBS. Post-wash, cell permeabilization was carried out with 0.5% PBST (Triton X-100 in PBS) at RT for 20 minutes. Click-iT reaction cocktail was prepared according to the composition given below.

| Click-iT™ Plus EdU Cell Proliferation Kit | Click-iT™ EdU Imaging Kit |
| --- | --- |
| 1X Click-iT reaction buffer – 880 µL<br>Copper protectant – 20 µL | Add the following amounts of reagents to the given amber vial in the kit containing |
| • Alexa Fluor picolyl azide (Alexa Fluor 488 used here) – 2.5 µL<br>• 1X Click-iT EdU buffer additive – 100 µL | Alexa Fluor picolyl azide (Alexa Fluor 488/594/647 azide were used here)<br>• 1X Click-iT reaction buffer – 100 µL<br>• CuSO<sub>4</sub> – 800 µL<br>• 1X Click-iT EdU buffer additive – 100 µL |

Permeabilization buffer was removed, and the cells were washed twice with 3% BSA in PBS. Cells were then incubated in the Click-iT reaction cocktail prepared, for 30 mins at RT, followed by a wash with 3% BSA in PBS. Lastly, the samples were counterstained for cell nuclei using DAPI (1μg ml^-1^ in PBS, Invitrogen) before imaging using widefield microscopy.

### Widefield microscopy

Widefield fluorescence images were acquired using 20X air objective (HC PL FLUOTAR L PH1 20x, NA=0.4, Leica) mounted on a Leica DMi8 microscope. Images were acquired using a Leica DFC9000 GTC sCMOS camera. Time-lapse images were taken inside a stage-top live chamber, maintained at 37°C and 5% CO2.

### Confocal microscopy

Fluorescence images were acquired using a 60X oil objective (PlanApo N 60x Oil, NA=1.42, Olympus) mounted on an Olympus IX83 inverted microscope equipped with a scanning laser confocal head (Olympus FV3000). Time-lapse images of live samples were done in the live-cell chamber provided with the microscopy setup. 25 mM HEPES buffer (Gibco) was used to maintain pH.

### Inhibition studies

For all inhibition studies, cells in glass bottom dishes were incubated with the required concentration of the inhibitor in DMEM at 37°C in a 5% CO2 humidified incubator before imaging. The cells were treated with FCCP (Sigma, 5μM) - a mitochondrial potential decoupler - for 2mins, to short-circuit the potential across the mitochondrial membrane. Myosin II inhibitor blebbistatin (Sigma, B0560, 30 mins treatment, 50 μM) and ROCK inhibitor Y27632 (sigma, 30 mins treatment, 30 μM), were used to disrupt the actomyosin contractility. All the drugs were dissolved in DMSO to prepare the primary stock solution.

### System biology modelling and simulation

The model can be described as follows:

- Grid system (100X100), each grid can have maximum one cell
- Simulation starts 500 cells randomly placed across the whole grid.
- Each cell has 2 key properties: activation level (mitochondrial activity) and proliferation/division time.
- Both the properties depend sigmoidally on the local density of the cell, calculated as the fraction of the occupied grids in a certain radius

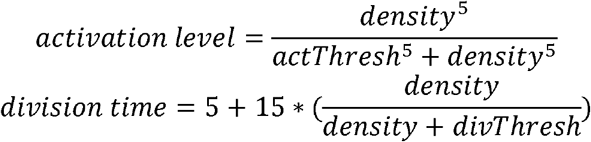

The activation level has been made more sensitive to the density than the division time by the use of the exponent (5) in its equation, owing to the fact that metabolic reactions react faster to the environmental cues than proliferation event.
- The density for each cell is calculated as the fraction of cells present in a 5X5 square around the cell. In other words, each cell is sensitive to a distance of 2 grids around it, represented as the R neighbourhood where R = 2.
- Each cell has a lifetime. Once the lifetime crosses the division time of the cell, the cell can divide if there is an empty grid in its R neighbourhood.
- At each time step, all cells are updated in three ways: their activation level changes based on the local density, their division time is updated based on the local density and they divide if the conditions specified previously are met.

### Image analysis

Image analyses were done by Fiji, custom written MATLAB scripts, Tissue Analyzer and cellpose [37]. Custom written MATLAB script was used for the TMRM intensity collectivity related analysis in Fig 1. Also, the geometrical parameters – local density, area, cell shape index, aspect ratio and TMRM intensity collectivity related analysis were carried out using custom written MATLAB scripts. All the cell-boundary segmentation was carried out either using custom written MATLAB code or using cellpose.

A custom written MATLAB code was used to calculate YAP Nuclear/Cytoplasmic ratio. Nuclear/Cytoplasmic ratio of YAP is defined as

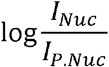

Where *I_Nuc_* is the average YAP intensity in the cell nucleus and *I_P.Nuc_* is the average YAP intensity in the perinuclear region [38]. Nucleus for each was segmented from the DAPI stained fluorescence image using cellpose. The perinuclear region is the dilated region of nuclear mask by a structural element of size 2. The cell boundaries were calculated by Voronoi tessellation of the cell centroids.

### Bayesian force inference

Bayesian force inference was implemented in a custom MATLAB script. The mathematical formulation of the method was first introduced by Ishihara and coworkers [29].

### Statistical analysis

Statistical analyses were carried out in GraphPad Prism 9. Statistical significance was calculated by *paired t-test with Welch’s correction or Wilcoxon matched-pairs signed rank test* as mentioned in the corresponding figure. Bar-whisker plots were displayed as mean ± max and min values. p-values greater than 0.05 were considered to be statistically not significant. No statistical methods were used to set the sample size. Quantification was done using data from at least three independent biological replicates. For analysis involving live-imaging experiments, data were collected from at least three independent experiments. All the experiments with representative images were repeated at least thrice.

## Supporting information

Combined Supplementary Information

## AUTHOR CONTRIBUTIONS

B.T., R.M., and S.K. performed the experiments. K.H. performed the model simulation. B.T., S.K., and T.D. conceived the project. T.D. supervised the experimental part while M.K.J. supervised the modelling part. B.T., K.H., M.K.J., and T.D. wrote the manuscript. All authors commented on the manuscript.

## ACKNOWLEDGEMENT

We thank the Collective Cellular Dynamics (CCD) laboratory members for critical discussions, Raphaël Clément for the source code of the Bayesian force inference program, and Manish Jaiswal and (late) Surajit Sengupta for their comments and suggestions. T.D. is a DBT/Wellcome Trust India Alliance intermediate fellow and partner group leader of the Max Planck Society (MPG), Germany. This work is funded by DBT/Wellcome Trust India Alliance (grant no. IA/I/17/1/503095 to T.D.). We also acknowledge the intramural funds at Tata Institute of Fundamental Research, Hyderabad from the Department of Atomic Energy Energy, India (under Project Identification No. RTI 4007) and the generous funding by the Human Frontier Science Program (HFSP).

