## Supplementary material for "Collective heterogeneity of mitochondrial potential in contact inhibition of proliferation": Combined Supplementary Information

Thurakkal *et al*.

This document contains:

1. Supplementary Table 1: Antibody information
2. Supplementary Figure Legends
3. Supplementary Video Legends

**Supplementary Table 1. Antibody information**

| Antibodies/Fluorophore/Dye | Catalog No. | Manufacturer | Dilution/Working concentration |
|---|---|---|---|
| TMRM | I34361 | Invitrogen | 100nM |
| JC-1 | T3168 | Invitrogen | 2μg/ml |
| DAPI (4′,6-diamidino-2-phenylindole) | D1306 | Invitrogen | 1 μg $ml^{-1}$ |
| YAP/TAZ (D24E4) Rabbit mAb | 8418 | CST | 1:100 |
| Tom20 (D8T4N) Rabbit mAb | 42406 | CST | 1:100 |
| Alexa Fluor 488, Goat anti-Rabbit IgG (H+L) Cross-Adsorbed Secondary Antibody | A-11008 | Invitrogen | Same as per primary antibody dilution |

-----------------------------------------------------------------------------------------------------------

**Supplementary Figure Legends**

**Supp. Figure 1: Metabolism shows different patterns of collectivity at different stages of CIP. a)** Scatter plot showing the change in the fraction of EdU-positive cells for different stages of CIP. **b)** High-resolution confocal image of MDCK stained with TMRM. The yellow dashed lines show patches of cells with low $\Delta\Psi_M$. Scale bar is 50um **c)** Representative TMRM intensity distribution at different global density regimes. Left to right: Stage 1, 2 and 3. The red arrow points to the peak at the near-zero intensity in stage 2. **d)** JC-1-stained images of MDCK monolayer corresponding to the three different stages of CIP. From left to right: Stage 1, 2 and 3. An increase in the levels of JC-1 aggregates can be seen with the increase in density.

**Supp. Figure 2: Quantification of local density and Tom20 correlation with TMRM intensity. a)** *Left panel***:** Line plots showing the average TMRM intensity for the cell population at high local density regions (top 20%), for different sizes of unit neighborhood box sizes (shown in the x-axis). Box sizes shown are in the unit of average cell diameter. Each line corresponds to one sample *Right panel*: Line plots showing the average TMRM intensity for the cell population at low local density regions (bottom 20%), for different sizes of unit neighborhood box sizes (shown in the x-axis). Box sizes shown are in the unit of average cell diameter. Each line corresponds to one sample. **b)** Left panel shows the centroid positions of each cell in a monolayer. The yellow box represents the unit-area box used to calculate local density at each point. The middle and right panels show the heatmaps of the cell monolayer comparing the local density field and average TMRM intensity respectively. **c)** Box and whiskers plot showing Tom20 intensity in cells with low and high TMRM intensity. Statistical significance was assessed using the Wilcoxon matched-pairs signed rank test.

**Supp. Figure 3: Dynamic model. a)** Histogram for activation corresponding to T=10,40 and 65 (Representing 3 stages)

**Supp. Figure 4: YAP localization. a)** *Top panel*: Fluorescence image of MDCK cells immune-stained with YAP antibody at stages 1, 2 and 3 of CIP. *Bottom panel*: Heatmap showing the cell-wise nuclear-cytoplasmic ratio. The cell boundaries are found by Voronoi tessellation. Patterns of YAP nuclear/cytoplasmic localization change with an increase in cell density, like heterogeneity in $\Delta\Psi_M$. Scale bar is 100μm.

**Supp. Figure 5: Cell pressure is correlated with $\Delta\Psi_M$. a)** Heat map comparing TMRM intensity and cell pressure for MDCK cell monolayer.

-----------------------------------------------------------------------------------------------------------

**Supplementary Video Legends**

**Supp. Video 1: Simulation dynamics for different activation and division thresholds. a)** Simulated dynamics of the system for an activation threshold of 0.9 and division threshold of 0.3. **b)** Simulated dynamics at high division threshold (0.9). The activation threshold is 0.9. **c)** Simulated dynamics at low activation threshold (0.3). The activation threshold is 0.3.

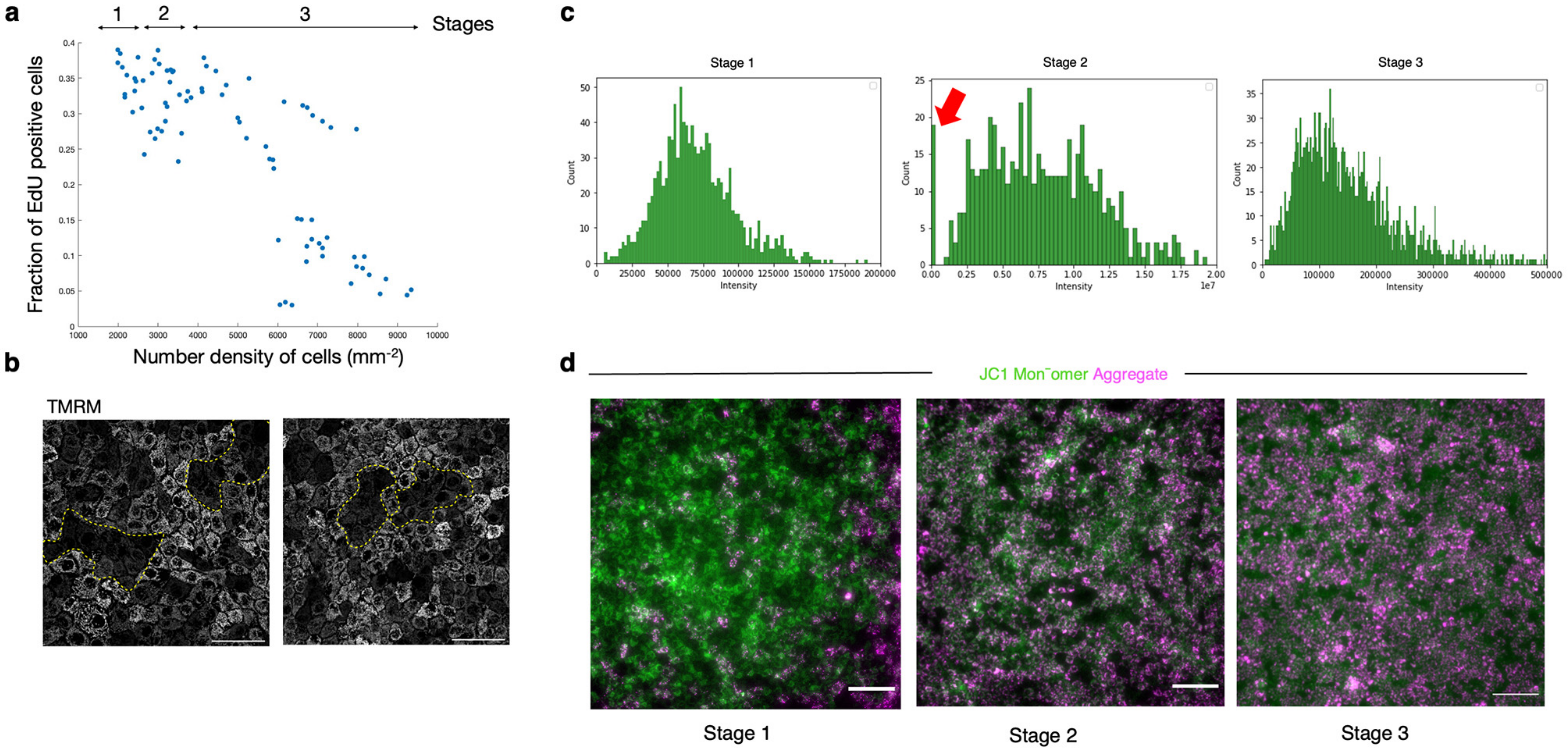


**Supplementary Figure 1**

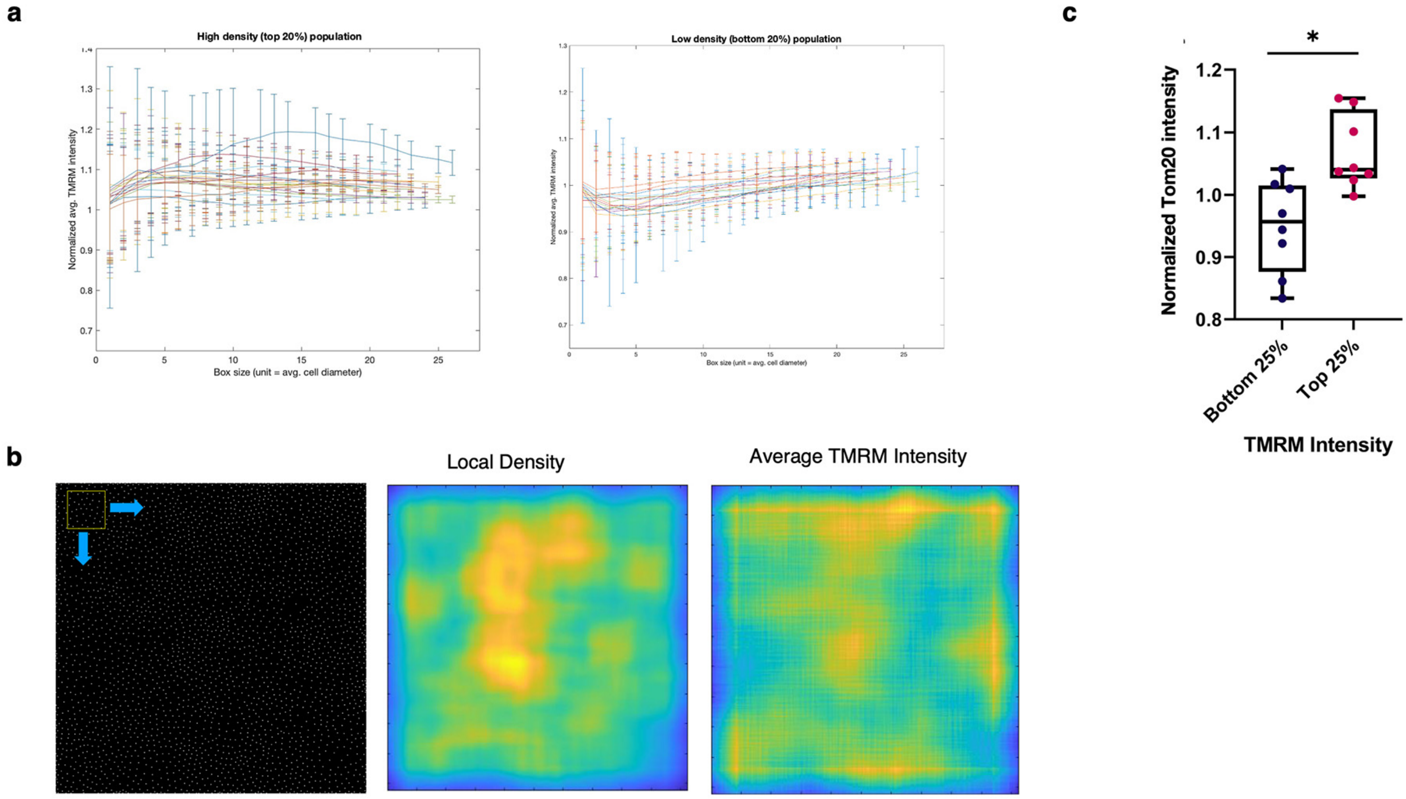


Supplementary Figure 2

a

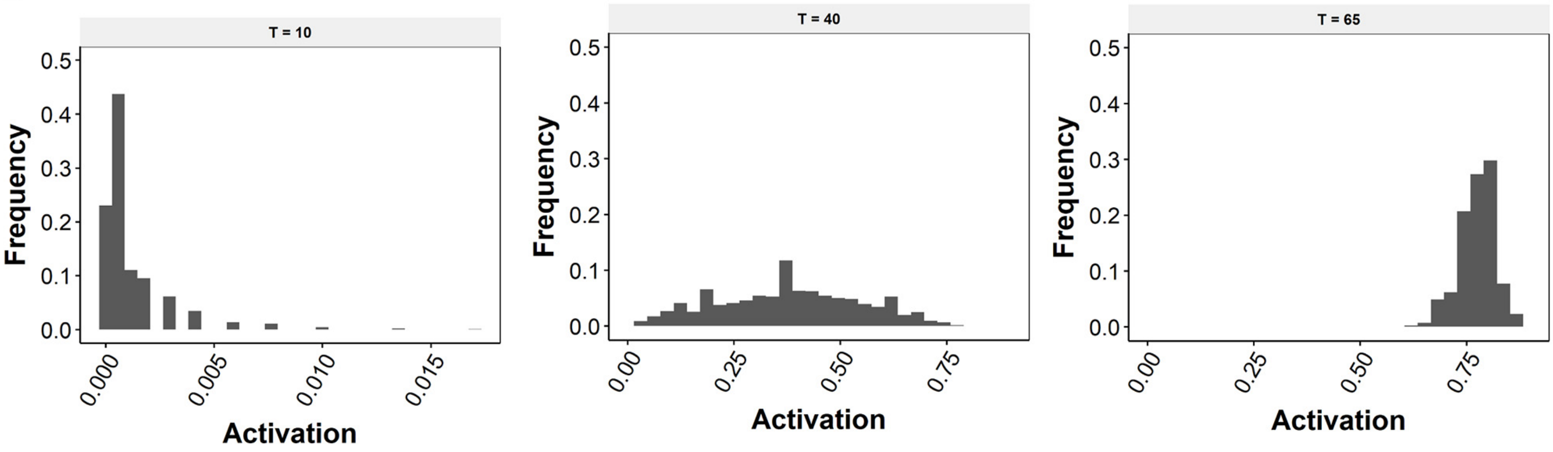


**Supplementary Figure 3**

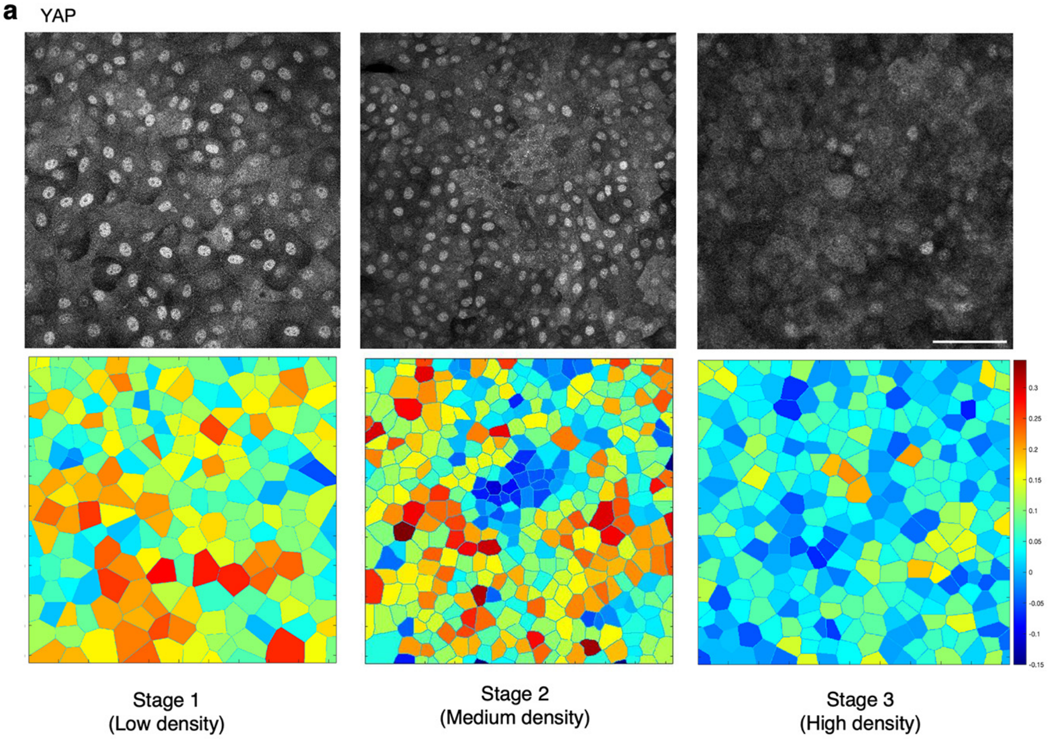


**Supplementary Figure 4**

**a**

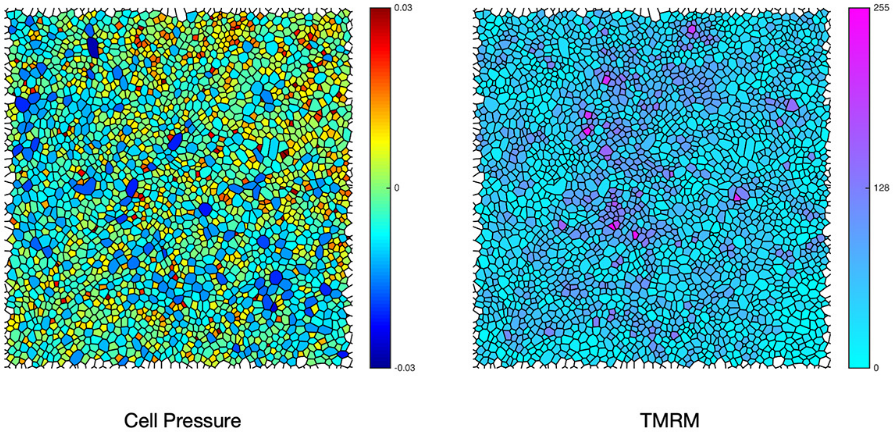


**Supplementary Figure 5**
